# Plasma proteome signature of canine acute haemorrhagic diarrhoea syndrome (AHDS)

**DOI:** 10.1101/2023.11.03.565505

**Authors:** Lukas Huber, Benno Kuropka, Pavlos G. Doulidis, Elisabeth Baszler, Lukas Martin, Anda Rosu, Lisa Kulmer, Carolina Frizzo Ramos, Alexandro Rodríguez-Rojas, Iwan A. Burgener

## Abstract

Acute haemorrhagic diarrhoea is a common complaint in dogs. In addition to causes like intestinal parasites, dietary indiscretion, intestinal foreign bodies, canine parvovirus infection, or hypoadrenocorticism, acute haemorrhagic diarrhoea syndrome (AHDS) is an important and sometimes life-threatening differential diagnosis. There is some evidence supporting the link between *Clostridium perfringens* toxins and AHDS. These toxins may be partially responsible for the epithelial cell injury, but the pathogenesis of AHDS is still not fully understood. Recent studies have suggested that severe damage to the intestinal mucosa and associated barrier dysfunction can trigger chronic gastrointestinal illnesses. Besides bloodwork and classical markers for AHDS such as protein loss and intestinal bacterial dysbiosis, we focused mainly on the plasma-proteome to identify systemic pathological alterations during this disease and searched for potential biomarkers to improve the diagnosis. To accomplish the goals, we used liquid chromatography-mass spectrometry. We compared the proteomic profiles of 20 dogs with AHDS to 20 age-, breed-, and sex-matched control dogs. All dogs were examined, and several blood work parameters were determined and compared, including plasma biochemistry and cell counts. We identified and quantified 207 plasmatic proteins, from which dozens showed significantly altered levels in AHDS. Serpina3, Lipopolysaccharide-binding protein, several Ig-like domain-containing proteins, Glyceraldehyde-3-phosphate dehydrogenase and Serum amyloid A were more abundant in plasma from AHDS affected dogs. In contrast, other proteins such as Paraoxonase, Selenoprotein, Amine oxidases, and Apolipoprotein C-IV were significantly less abundant. Many of the identified and quantified proteins are known to be associated with inflammation. Other proteins like Serpina3 and RPLP1 have a relevant role in oncogenesis. Some proteins and their roles have not yet been described in dogs with diarrhoea. Our study opens new avenues that could contribute to the understanding of the aetiology and pathophysiology of AHDS.

## Introduction

Acute Haemorrhagic Diarrhoea Syndrome (AHDS) in dogs is a potentially life-threatening gastrointestinal disorder with a high prevalence [1–3]. Characterised by a sudden onset of bloody diarrhoea, vomiting, and profound dehydration resulting in haemoconcentration, AHDS treatment poses significant challenges for veterinary practitioners [4]. The condition can progress rapidly, leading to severe complications and, in some small number of cases, the outcome can be fatal if not adequately addressed [1]. Persistent vomiting and profuse diarrhoea lead to rapid bodily fluid loss, severe dehydration, and an electrolyte imbalance [5]. Dehydration affects the dog’s overall health, leading to weakness, lethargy, and potential organ dysfunction. Blood loss can lead to anaemia and contribute to morbidity [6]. In severe cases, AHDS can trigger systemic inflammatory response syndrome (SIRS), an aggressive immune reaction that causes widespread inflammation. This reaction can progress to sepsis or even septic shock, leading to a life-threatening drop in blood pressure and organ dysfunction [7]. Septic shock requires immediate and intensive medical intervention to decrease the risk of death but is, in any case, associated with a high mortality [8]. Despite its prevalence and numerous publications investigating this disease, the aetiology of AHDS is unknown. Several pathogens are associated with AHDS in dogs, including some viruses, bacteria, and parasites. In AHDS, significant damage to the lining of the stomach and intestines occurs. It is suspected that the involved pathogens could directly attack and destroy the intestinal cell lining, leading to haemorrhage and ulceration [8]. Some strains of *Clostridium perfringens*, a bacterium that is an ordinary member of the canine microbiome, produce the pore-forming toxins NetE and NetF, which are commonly discussed in the literature as a major cause of gastrointestinal epithelial necrosis leading to AHDS [9]. As the incidence of AHDS is next to parasitic diseases and alimentary indiscretion probably the highest among acute gastrointestinal diseases in dogs, understanding the underlying aetiology and pathophysiology, identifying predisposing factors and developing effective management strategies are critical in mitigating the impact of this syndrome on canine health [10]. While MS-based proteomic analysis in veterinary medicine is less extensive than in human clinical research, notable studies have focused on samples from dogs afflicted with diverse diseases [11]. The application of mass spectrometry for quantitative proteome analysis in veterinary medicine holds significant promise for enhancing our understanding of the underlying mechanisms of various diseases like AHDS in dogs. These techniques enable researchers to identify and quantify proteins with a higher degree of precision, aiding in discovering of potential biomarkers and therapeutic targets. Although the current body of research is comparatively limited, the growing attention to this cutting-edge approach in veterinary science is encouraging [12].

This article reports a quantitative proteomic study of canine plasma from patients affected by AHDS and its comparison with plasma from healthy controls. The main goal of this study is to describe the general perturbation of the blood induced by AHDS and the exploration of new potential biomarkers that could help to understand the pathogenesis of AHDS better, improve future diagnostics and aid in the treatment and follow-up of the disease.

## Results and Discussion

Breeds represented in this study in the AHDS group included Crossbreed (4), Chihuahua (3), Maltese (2), Pomeranian (2), Yorkshire Terrier (2), Chinese Crested (1), English Bulldog (1), Golden Retriever (1), Havanese (1), Labrador Retriever (1), Poodle (1) and Schnauzer (1). The control group included Crossbreed (8), Poodle (3), German Shepherd Dog (2), English Toy Terrier (1), French Bulldog (1), Golden Retriever (1), Husky (1), Labrador (1), Pomeranian (1) and Västgötaspets (1).

The median age of the dogs in the study group was 3.5 years (1–11 years), and the age of the dogs in the control group was 4.5 years (1-10 years). The mean body weight was 5.3kg (1.5-27.5kg), while the control group weighed 15.9 kg (3.5-33kg). We included 12 female (6 intact and 6 neutered) and 8 male (5 neutered and 3 intact) dogs in the AHDS group. In the control group 14 female (9 intact and 5 neutered) and 6 male (1 intact and 5 neutered) dogs were examined.

All the dogs in the control group were regarded as clinically healthy and showed no abnormalities at physical examination. Of the AHDS dogs the most common reported presenting complains besides diarrhoea were vomiting (75%, 15/20), apathy (45%, 9/20) and anorexia (45%, 9/20) and the most frequent clinical signs were tachycardia (>100beats/min) (85%, 17/20) and abdominal discomfort on palpation (85%, 17/20). Complete blood count and a plasma biochemistry panel containing glucose level, urea, creatinine, total proteins (Tp), albumin, ALT, and ALKP, were carried out for all dogs except one from the AHDS group.

A comparative analysis of blood cell counts between the AHDS and control groups revealed significant differences in various markers (Figure 1). The AHDS group exhibited a significantly lower mean corpuscular volume (MCV) with *p*=0.0019 along with reduced lymphocyte counts (*p* = 0.0013) and eosinophil counts (*p*=0.013). In contrast, sickened dogs demonstrated a higher abundance of leukocytes (*p*=0.00024), monocytes (*p*=0.0061), and neutrophils (*p*=0.0009) than the control group. The rest of the blood cell markers were non-significant (see Figure S1).

**Figure 1.**
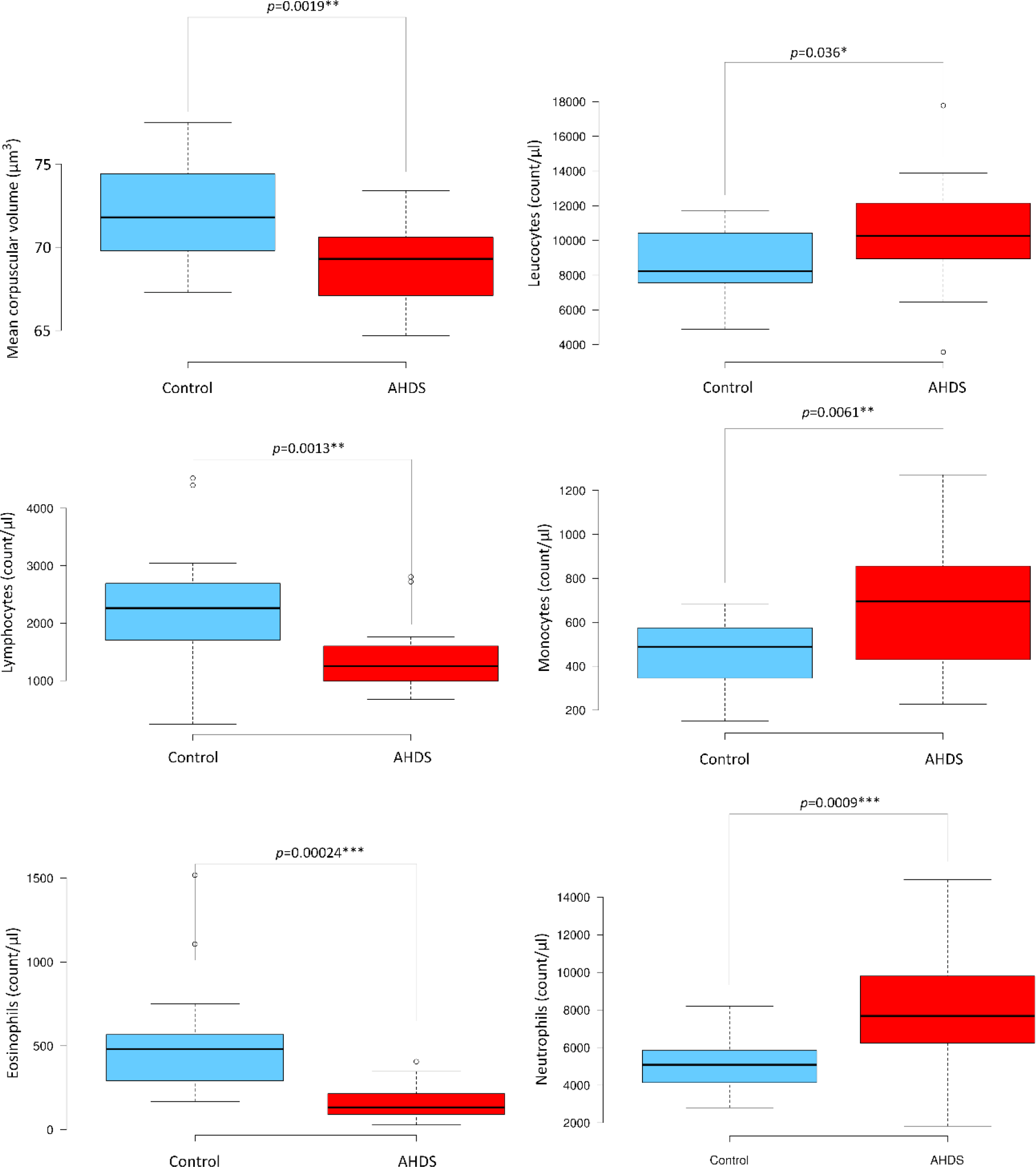
Boxplot analysis of significant blood cell markers in the AHDS and control groups. This figure presents boxplots showcasing the significant variations in blood cell markers observed between the AHDS and control groups. Statistical differences were determined using either the student’s t-test for samples with equal variance or the Welch test for samples exhibiting differing variances. Significance levels were established as *p* < 0.05, significant *; *p* < 0.01**, highly significant; and *p* < 0.001*\*\**, very significant.

Like the changes in the complete blood cell count, also the biochemistry panel shared significant differences between the AHDS cohort and healthy controls. AHDS patients displayed a notable decrease in the abundance of total proteins (TP, *p*=<0.001), also corroborated by lower levels of albumin (*p*=0.00022). Conversely, AHDS dogs exhibited elevated levels of glucose (*p*= 0.0019) and alkaline phosphatase (*p*= 0.033). These findings are summarised in Figure 2, while the results for other plasma biomarkers can be found in Figure S2.

**Figure 2.**
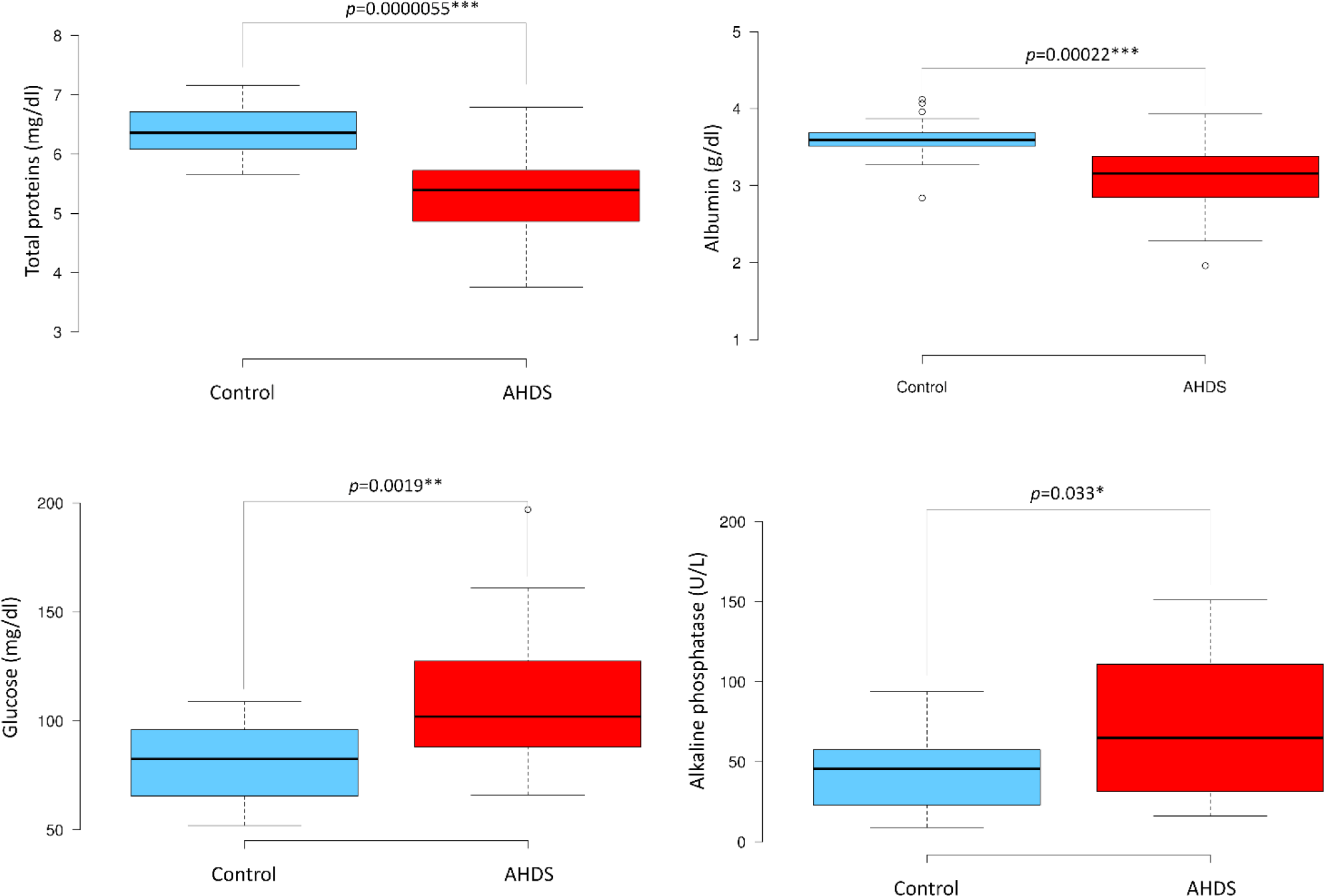
Boxplot analysis of significant biochemistry parameters in the AHDS and control groups. This figure provides a boxplot analysis depicting significant variations in select plasma biochemistry markers between the AHDS and control groups. Statistical differences were assessed using either the student’s t-test for samples with equal variance or the Welch test for samples displaying varying variances. Significance levels were established as *p* < 0.05, significant *; *p* < 0.01**, highly significant; and *p* < 0.001*\*\*\**, extremely significant.

Next, we used label-free quantification mass spectrometry to analyse canine plasma from patients affected AHDS in comparison with healthy controls. This study aims to uncover average traits or typical patterns in AHDS with the potential for discovering new biomarkers. The experiment was conducted using pooled samples, comprising ten independent plasma pools (five from AHDS samples and five from control samples), each collected from four randomly selected individuals.

A comprehensive examination of the data revealed that the pools from AHDS patients exhibited a distinct proteome signature compared to those from healthy dogs. Principal component analysis of all quantified proteins from both groups demonstrated the segregation of samples into two separate clusters, indicating the sensitivity of the proteomic analysis in capturing these differences (see Figure 3A).

**Figure 3.**
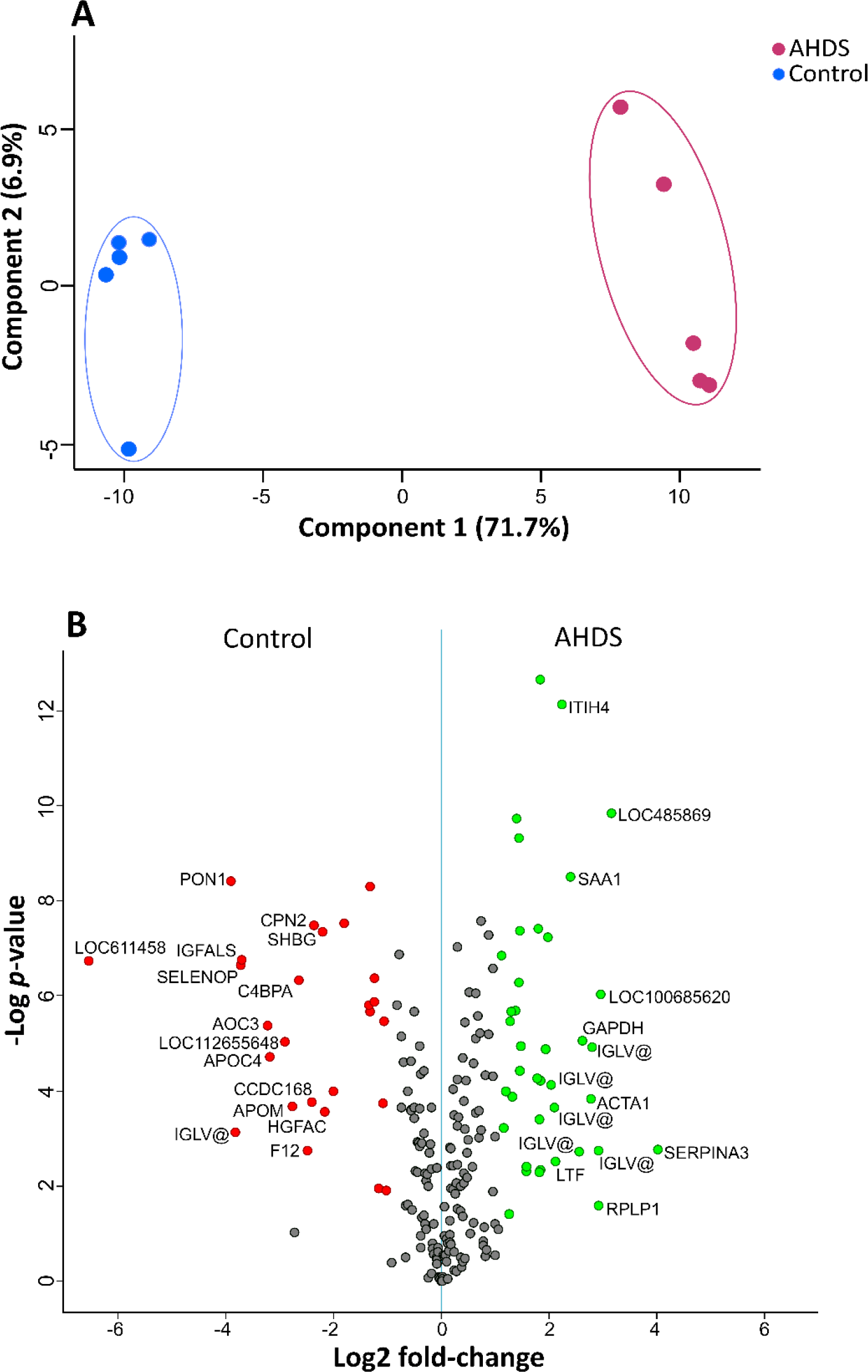
(A) Principal Component Analysis (PCA) of quantitative proteome analysis reveals the distinct separation between healthy dogs and those affected by AHDS. (B) Volcano plots were generated to compare plasma pools from healthy dogs with those from five AHDS-affected individuals (n=4 each). Gene names were employed instead of protein names. Upregulated proteins are denoted by green dots, signifying higher abundance in AHDS dogs compared to healthy ones (p-value < 0.05 and a log2 fold-change greater than 2). Conversely, downregulated proteins are indicated by red dots, indicating lower abundance in AHDS group compared to the control (*p*-value < 0.05 and a log2 fold-change smaller than −2). Non-significantly differentially expressed proteins are depicted as grey dots. A comprehensive list of identified and quantified proteins is available in Table S1.

Among the 207 quantified proteins, we identified 95 proteins exhibited elevated levels, and 67 showed diminished abundance in the plasma of AHDS-affected individuals compared to healthy dogs, highlighting the discernible impact of the disease (Figure 3B, Table S1). Considering these findings, our focus is directed toward proteins that demonstrated the most significant increments or reductions associated with AHDS.

Notably, within the group of proteins increased in AHDS, the following top 8 proteins stand out: SERPINA3, Lipopolysaccharide-binding protein (encoded by the locus LOC485869), Ribosomal protein lateral stalk subunit P1 (RPLP1), Actin alpha 1 of skeletal muscle (ACTA1), Glyceraldehyde-3-phosphate dehydrogenase (GAPDH), Serum amyloid A (SAA1), Inter-alpha-trypsin inhibitor heavy chain 4 (ITIH4) and Lactotransferrin (LTF).

Notably, a distinct case exists within our dataset - the variable region of lambda light chains, a pivotal component within all immunoglobulins, exhibited altered levels in AHDS samples compared to the control group. These proteins exhibit notable diversity, and in our study, we identified 14 distinct protein variants showing higher abundance in AHDS (see Table S1). In a converse trend, two variants experienced downregulation. These specific protein groupings have been denoted as IGLV@, as referenced in specific databases [13].

Other proteins that exhibited significantly increased abundance in AHDS include Serum amyloid A (SAA1), Inter-alpha-trypsin inhibitor heavy chain 4 (ITIH4), and Lactotransferrin (LTF). Proteins with a significantly increased abundance in the control group, or conversely, a lower abundance in the AHDS group, comprised Pregnancy zone protein-like (encoded by the gene LOC611458), Paraoxonase (PON1), Selenoprotein P (SELENOP), Insulin-like growth factor binding protein acid labile (IGFALS), and two Amine oxidases (AOC3 and LOC112655648). Other proteins that were depleted in the AHDS group involved Apolipoprotein M (APOM), complement component 4 binding protein alpha (C4BPA), and coagulation factor XII (F12), Carboxypeptidase N (CPN2), Sex hormone binding globulin (SHBG), and the HGF activator (HGFAC).

Using the dataset of quantified proteins, we conducted gene ontology enrichment analysis [14] to investigate the functional significance of differentially expressed proteins (DEPs) in AHDS patients compared to the healthy control group (Figure 4). The analysis highlighted some homologous proteins related to Alzheimer’s disease in humans, as well as Parkinson’s and Huntington’s diseases. Additionally, notable protein classes in both upregulated and downregulated groups included coagulation-related proteins such as those involved in the plasminogen activation cascade and inflammation-related proteins.

**Figure 4.**
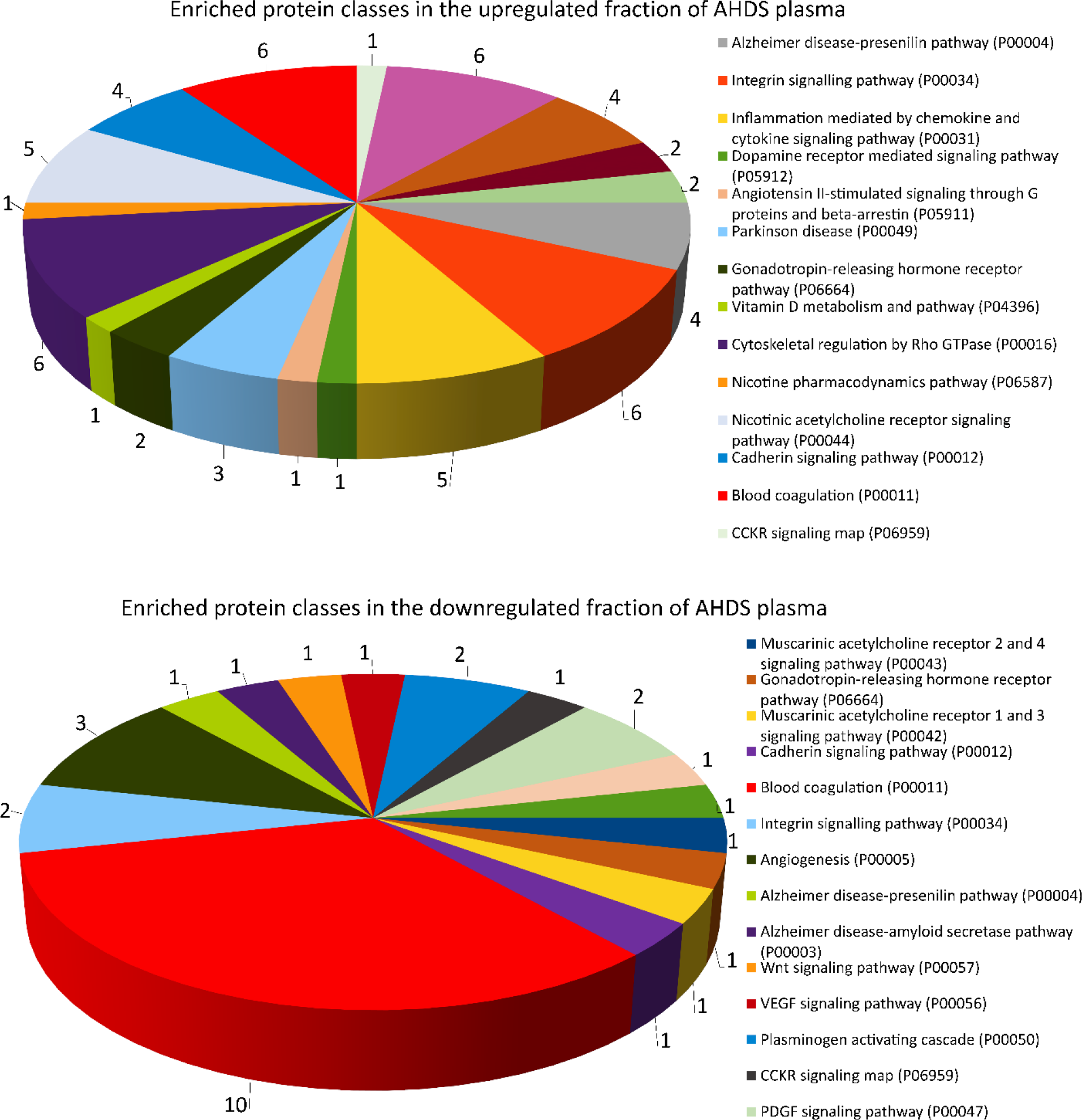
The figure illustrates the Gene Ontology (GO) analysis results conducted on the plasma proteome of dogs affected by AHDS. The analysis categorized the proteins according to their pathways using Panther software [14]. Each slice represents the set of genes (hits) for each specific functional category or pathway. Only genes that had a functional category described are represented.

We have utilised a network analysis rooted in protein-protein interactions, drawing upon functional insights sourced from the STRING database, to visually depict the overarching protein alterations in the plasma of ADHS patients. This thorough examination encompasses both direct (physical) and indirect (functional) associations among proteins [15]. To elucidate the impact of AHDS on plasmatic protein levels, we superimposed our proteomic datasets onto the known protein-protein interaction database for dogs. Notably, the proteins exhibiting up-regulation and down-regulation in AHDS displayed a remarkable degree of interconnectivity (Figure 5). This striking observation hints at extensive proteome-wide adjustments in response to the pathological condition, offering insight into the modifications within various protein clusters.

**Figure 5.**
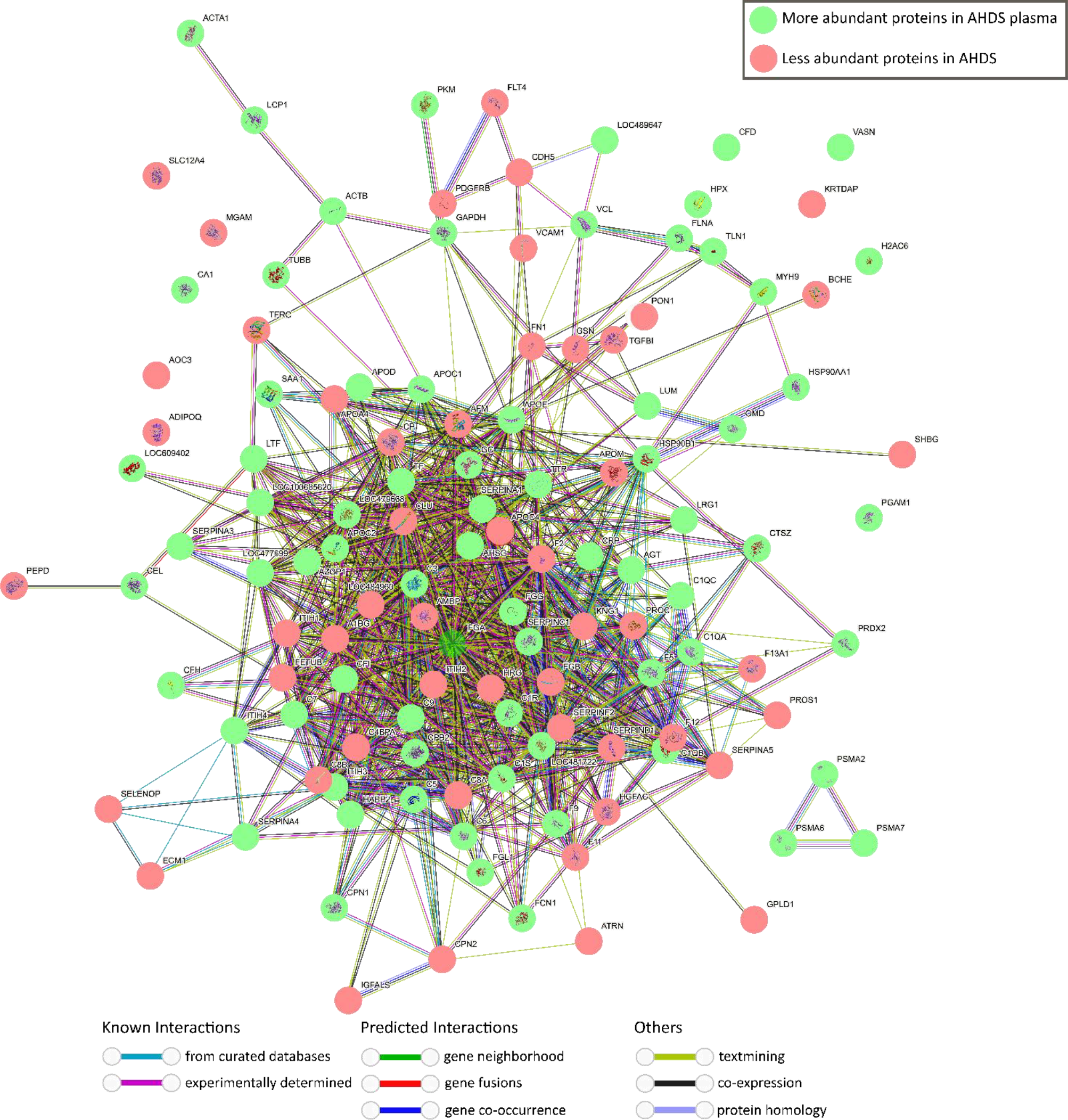
Network visualization of proteins overrepresented (green) and depleted (red) in plasma of dogs with AHDS compared to the plasma of healthy dogs using STRING database [33]. The network diagram depicts the interactions between various canine plasma proteins using as inclusion a significant FDR-adjusted *p*-value regardless of the level of expression. Each node represents a unique protein, and the edges connecting nodes indicate known or predicted interactions. Node colour indicates the type of interaction evidence, ranging from experimental data to curated databases and text mining.

Collecting intestinal tissue samples from animals can be challenging in most cases for ethical or clinical issues. However, when it comes to investigating the development of diarrhoea in animals, plasma emerges as an optimal choice for sampling. This preference arises from its comparatively stable chemical and physical properties, rich protein content, and relatively reliable collection techniques [16]. We can add to these facts that since haemorrhagic diarrhoea affects the overall physiological functioning of the body, plasma protein profiling offers valuable insights into these perturbations, making it a very suitable material for diagnosis and biomarker discovery.

It is important to note that in cases of intestinal blood loss, patients experience a depletion of electrolytes and water, causing haemoconcentration, accompanied by the loss of specific proteins that impact the whole body’s homeostasis [17]. This explains why the total protein and albumin levels in AHDS dogs are lower compared to the control group although they are haemoconcentrated. These haemoconcentration and selective protein losses trigger a reactive response that in turn affects the levels of other proteins. As a result, the findings presented here may reflect a delicate trade-off among these factors and the overall homeostatic disruption observed in AHDS patients.

The classical clinical blood analysis revealed significant imbalances in blood cell counts and plasma biochemistry. The discrepancies can be partially attributed to factors such as intestinal blood loss and electrolyte imbalances, which result in haemoconcentration [4]. Neutrophilia and monocytosis as well as a rise in blood glucose fit to the picture of a stress leukogram, triggered by the effects of adrenalin and cortisol during acute illness, but can also be seen in systemic inflammation in general. In some instances, these factors may lead to alterations that either overestimate or underestimate the quantification of specific biomarkers. This phenomenon arises from the complex interplay of various mechanisms within the organism. For instance, the body may attempt to replenish specific blood components when possible [18]. At the same time, alterations in other parameters can relate to immune responses or the expenditure of proteins, such as coagulation factors.

Some of the altered proteins found in the ADHS samples are already known to be involved in other gastrointestinal disorders. For instance, an increase of the amine oxidases (AOC3 and AOX2) is considered an indicator of intestinal injury caused by some anti-cancer drug treatments [19–21].

Another good example is the protein SERPINA3, the protein which showed the strongest increase in the ADHS plasma samples compared to healthy controls. SERPINA3, also known as Alpha-1-antichymotrypsin (ACT), belongs to the serine protease inhibitor (serpin) family. Its primary function is to regulate and inhibit the activity of certain proteases, specifically chymotrypsin-like enzymes. SERPINA3 acts as an inhibitor of chymotrypsin-like proteases. By binding to these proteases and forming a stable complex, SERPINA3 helps to prevent excessive proteolysis, which is essential for maintaining proper protein balance and preventing tissue damage [22]. In some inflammatory processes, SERPINA3 is also implicated in the modulation of certain components of the immune system and cytokines, affecting immune responses and inflammation. Changes in SERPINA3 levels have been associated with various inflammatory conditions and diseases. SERPINA3 may also play a role in regulating fibrinolysis, the breaking down of blood clots. It interacts with plasmin, an enzyme involved in fibrinolysis, and can influence the balance between clot formation and clot dissolution [23]. SERPINA3 is also involved in tissue repair and remodelling, modulating tissue responses to injury, and promoting healing. Some studies have pointed out that SERPINA3 levels may be altered in specific cancer types and neurodegenerative diseases [22].

Another highly elevated protein is the RPLP1, which belongs to the ribosomal P complex, which consists of the acidic ribosomal P proteins. The presence of this intracellular protein in the bloodstream may serve as an indicator of significant tissue damage and the release of cellular contents into the circulatory system in AHDS affected dogs. RPLP1 overexpression not only immortalises primary cells but also plays a significant role in their transformation. Furthermore, there is evidence of RPLP1 protein being upregulated in certain human cancers [24]. In the same direction, it is known that Glyceraldehyde-3-phosphate dehydrogenase, another upregulated protein in AHDS, is overexpressed in colorectal cancer onset [25]. These findings could be another link between gastrointestinal affection like AHDS and the development of future tumours, which is another reason to enhance better understanding and management of the disease.

It is also intriguing to speculate about the relationship between some of the protein classes that exhibited altered levels in this study and their roles in conditions associated with the brain, including Alzheimer’s disease, Parkinson’s disease, and Huntington’s disease as shown here with the gene ontology analysis. For instance, older dogs often develop canine cognitive dysfunction, a condition that shares many similarities with Alzheimer’s disease. Dogs exhibit cognitive deficits that can be compared to human symptoms, such as disorientation, memory loss, changes in behaviour, and the presence of beta-amyloid plaques, which are commonly detected in both the extracellular space as senile plaques and around blood vessels in their brains. [26]. Various clinical, epidemiological, and immunological findings provide strong indications that the gut-brain axis is significantly and deeply impacted by alterations of gut microbiome [27].

Managing diarrhoea in dogs, whether related to conditions like AHDS or similar ailments, holds significance for the well-being of animal companions and human health, as coined by the concept of One Health [28]. Certain microorganisms found in these faecal deposits could serve as sources of zoonotic transmission for parasites such as *Giardia* or potential bacterial pathogens like *Campylobacter, E. coli*, and *Salmonella*. Hence, understanding and controlling diarrhoea in dogs is crucial for safeguarding both animal and human health.

The study’s reliance on a relatively small cohort of just 40 participants raises some limitations that must be considered when interpreting the results. The sample size may not provide a comprehensive representation of the broader population, limiting the generalizability of the findings. Furthermore, an additional challenge arises from the noticeable differences in body weight between the groups within the study, and the breed are not exactly matching. These variations can introduce confounding variables that may obscure some effects of the factors under investigation. Differences in breed and body weight can influence the outcomes in ways that are unrelated to our primary research question, potentially leading to skewed or biased results. Therefore, while the study provides valuable insights, these limitations should be considered when drawing conclusions or extrapolating the findings to a larger and more diverse population.

Our study opens several promising avenues for future research by uncovering the alteration of several proteins in plasma that were not previously reported in the literature for AHDS. First, we need to investigate further the role of the identified biomarker candidates in other diarrhoeic diseases to understand better if altered proteins are specific for AHDS or if changes in their expression levels also occur in other confounding conditions. A comparison with the proteomic profile in diseases like Parvovirosis or “inflammatory bowel disease” (IBD) would be a possible next step. Second, we should explore the connection between severe intestinal mucosal damage, barrier dysfunction, and the development of other chronic gastrointestinal diseases.

To improve the AHDS diagnostic accuracy and treatment strategies, future studies should focus on refining and expanding the pool of potential biomarkers we have identified here. Considering a broader range of canine populations and disease stages, additional studies are required to extend the validity of our results. Collaborative efforts to validate and translate these candidates into clinical practice could ultimately improve the prognosis and management of AHDS in dogs. Taken together, our research provides a foundation upon which future investigations can continue to advance our understanding and enhance the care of dogs affected by AHDS. To the best of our knowledge, this is the first study that employs a quantitative plasma proteomic approach to gain a deeper understanding of AHDS.

## Conclusions

In conclusion, acute haemorrhagic diarrhoea syndrome in dogs presents a significant clinical challenge, often leading to severe health complications. While several factors, including *Clostridium perfringens* toxins, have been implicated in its development, the exact pathogenesis of AHDS remains elusive. Our study took a comprehensive approach, exploring the proteomic profiles of dogs afflicted with AHDS and comparing them to reasonably well-matched control dogs. To the best of our knowledge, this is the first study that employs a quantitative plasma proteomic approach to gain a deeper understanding of AHDS. This in-depth analysis uncovered substantial alterations in the levels of numerous proteins present in plasma, shedding light on potential biomarkers for diagnostic and therapeutic advancements and possibly therapeutic interventions. SerpinA3, Lipopolysaccharide-binding protein, Ig-like domain-containing proteins, Glyceraldehyde-3-phosphate dehydrogenase, and Serum amyloid A emerged as candidates with increased abundance in AHDS-affected dogs. In contrast, others exhibited decreased levels, such as Paraoxonase, Selenoprotein, Amine oxidases, and Apolipoprotein C-IV.

## Materials and methods

### Animals and sample collection

All the samples used for this work were surplus materials from previous studies. The previous studies were approved by the Ethics Committee of the University of Veterinary Medicine Vienna and the Austrian Federal Ministry of Science and Research (BMBWF-68.205/0216-V/3b/2019, 2021-0.518.574), and methods for sample collection were carried out in accordance with relevant Austrian guidelines and regulations. In this work, we analysed the samples of 20 client-owned dogs presented to the Clinic for Small Animal Internal Medicine of the University of Veterinary Medicine Vienna, Austria, over a period of one year and diagnosed with AHDS according to current diagnostic criteria from a still ongoing prospective study. We compared them to a healthy control group matched according to age, sex and a phylogenetic tree. Data collected from these animals included age, sex, breed, body weight, physical examination findings and clinicopathological values. Further diet and a history of previous gastrointestinal diseases were especially investigated during the anamnesis. Only dogs with acute bloody diarrhoea for less than three days, with haemoconcentration with a haematocrit of ≥50% and without a known history of chronic gastrointestinal disorders were included in the study.

As a control group, we used clinically healthy client-owned dogs (N=20) with no signs of disease that were presented to the Clinic for Small Animal Internal Medicine over a six-month period for clinical examination and blood sampling to be enrolled in a behavioural study from the Interuniversity Messerli Institute of Research (Vienna, Austria). Inclusion followed a thorough history, physical examination, and blood sampling. The samples were analysed including complete blood count, plasma biochemical profile and electrolytes measurements. All samples showed no significant pathological alterations.

The samples from both groups, diagnosed with AHDS and the healthy control group, were grouped into five independent pool sets, each created using the “tidyverse” R package to generate randomisation and group selection as previously described [12]. The procedure involved assigning a number to each sample. Upon execution, the computer script generated five random lists, each containing four individuals. These lists were used to prepare the pools. Samples were pooled to limit the effect of individual samples.

## Quantitative proteome by label-free liquid chromatography–mass spectrometry (LC–MS)

Blood samples were collected in 2 ml Vacuette tubes with lithium heparin (clinical chemistry) 13×75 green cap-white ring PREMIUM (Greiner Bio-One GmbH, Bad Haller Str. 32, 4550, Austria) and the plasma was centrifugated at 2000 x g for 5 minutes and stored in −20°C. Five microliters of plasma from each of the 40 individuals was collected from the supernatant after a centrifugation step at 20 000 x g at 4 °C. For each experimental replicate, five samples containing the pooled plasma of four randomly selected individuals were prepared. From each plasma pool, 3 μl were transferred to a tube containing 120 μl urea denaturing buffer (6 M urea, 2 M thiourea, and 10 mM HEPES; pH 8.0). Proteins were reduced by adding 5 μl dithiothreitol (10 mM) and incubated for 30 min at room temperature. Then, the samples were alkylated by the addition of 5 μl iodoacetamide solution (55 mM) and incubated at room temperature for another 30 min in the dark. All samples were diluted by adding four volumes of ammonium bicarbonate buffer (50 mM) to reduce the urea concentration. Thereafter, samples were digested overnight at 37 °C after adding 1 μg of trypsin protease (Thermo Scientific, USA). Next day, the samples were acidified by adding a final concertation of 5% acetonitrile and 0.3% of TFA and subsequently desalted via C18 StageTips with Empore™ C18 Extraction Disks [29]. Peptides eluted from the StageTips were dried by vacuum centrifugation. Peptides were reconstituted in 80 µl of 0.05% TFA, 4% acetonitrile and 1 µl of each sample was applied to an Ultimate 3000 reversed-phase capillary nano liquid chromatography system connected to a Q Exactive HF mass spectrometer (Thermo Fisher Scientific). Samples were injected and concentrated on a PepMap100 C18 trap column (3 µm, 100Å, 75 µm inner diameter [i.d.] × 20mm, nanoViper; Thermo Scientific) equilibrated with 0.05% TFA in water. After switching the trap column inline, LC separations were performed on an Acclaim PepMap100 C18 capillary column (2 µm, 100Å, 75 µm i.d. × 500mm, nanoViper; Thermo Scientific) at an eluent flow rate of 300 nl/min. Mobile phase A contained 0.1% (v/v) formic acid in water, and mobile phase B contained 0.1% (v/v) formic acid and 80% (v/v) acetonitrile in water. The column was pre-equilibrated with 5% mobile phase B followed by an increase to 44% mobile phase B over 100 min. Mass spectra were acquired in a data-dependent mode, utilizing a single MS survey scan (m/z 300–1650) with a resolution of 60,000, and MS/MS scans of the 15 most intense precursor ions with a resolution of 15,000. The dynamic exclusion time was set to 20 s, and the automatic gain control was set to 3 × 10^6^ and 1 × 10^5^ for MS and MS/MS scans, respectively. MS and MS/MS spectra were processed using the MaxQuant software package (version 2.0.3.0) with the implemented Andromeda peptide search engine [30]. Data were searched against the *Canis lupus familiaris* reference proteome (ID: UP000002254; downloaded from Uniprot.org on 29.03.2023; 59102 sequences) using the default parameters and enabling the options label-free quantification (LFQ) and match between runs. Data filtering and statistical analysis were conducted with the Perseus 1.6.14 software [31]. Only proteins that were identified and quantified with LFQ intensity values in at least three (out of five) replicates (within at least one of the two experimental groups) were used for downstream analysis. Missing values were replaced from normal distribution (imputation) using the default settings.

### Statistical analysis

Statistical differences in protein expression levels (LC-MS) were determined using Student’s t-tests with a permutation-based False Discovery Rate (FDR) of 0.05 implemented in the Perseus software [31]. Proteins with a minimum 2-fold intensity change compared to control (log2 fold-change ≥ 2 or log2-fold change ≤ −2) and a *p*-value ≤ 0.05 (FDR adjusted *p*-value) were considered significantly affected. To compare the blood markers, we employed Student’s t-tests to assess the significance of differences between two groups (AHDS and control) when variances were found to be equal through the Levene’s test. If variances were unequal, Welch’s test was utilised instead. All *p*-values less than or equal to 0.05 were considered significant. These test were performed using R computational language platform in R-studio version 2023.09.0+463 [32].

## Availability of data and materials

We have included all data generated or analysed during this comprehensive study within this published article and its supplementary information files.

## Authors’ contributions

Conceptualization: LH, ARR, and IAB. Methodology: LH, BK, PGD, ARR. Formal analysis: LH, BK, PGD, EB, LM, AR, LK, CFR, ARR, and IAB. Investigation: LH, BK, PGD, EB, LM, AR, LK, CFR, ARR, and IAB. Resources: BK, ARR, IB. Writing original draft: LH, ARR, PGD, IAB. Writing, review and editing: LH, BK, PGD, EB, LM, AR, LK, CFR, ARR, and IAB. Supervision: ARR and IAB. Project administration: ARR and IAB. Funding acquisition: ARR and IAB. Authorship: We declare that all the authors of this study have directly participated in the planning, execution, or analysis of the study, and all the authors have read and approved the final version submitted.

## Acknowledgements

For mass spectrometry, we would like to acknowledge the assistance of the Core Facility BioSupraMol supported by the Deutsche Forschungsgemeinschaft (DFG).

---

**Table S1.** Table S1 presents the results of the proteomic experiment analysis utilizing Perseus export, detailing the detection and quantification of plasma proteins in dogs affected by Acute Hemorrhagic Diarrhea Syndrome (AHDS) in comparison with a control group. The dataset encompasses five replicates from six distinct independent pools of plasma samples, comprising both AHDS-affected dogs and healthy controls. Statistical analysis was conducted employing the student t-test, and p-values were subjected to correction using the false discovery rate (FDR) methodology. The analysis was performed utilizing MaxQuant and Perseus software, employing label-free quantification of proteins through Liquid Chromatography-Mass Spectrometry (LC-MS).

**Figure S1.**
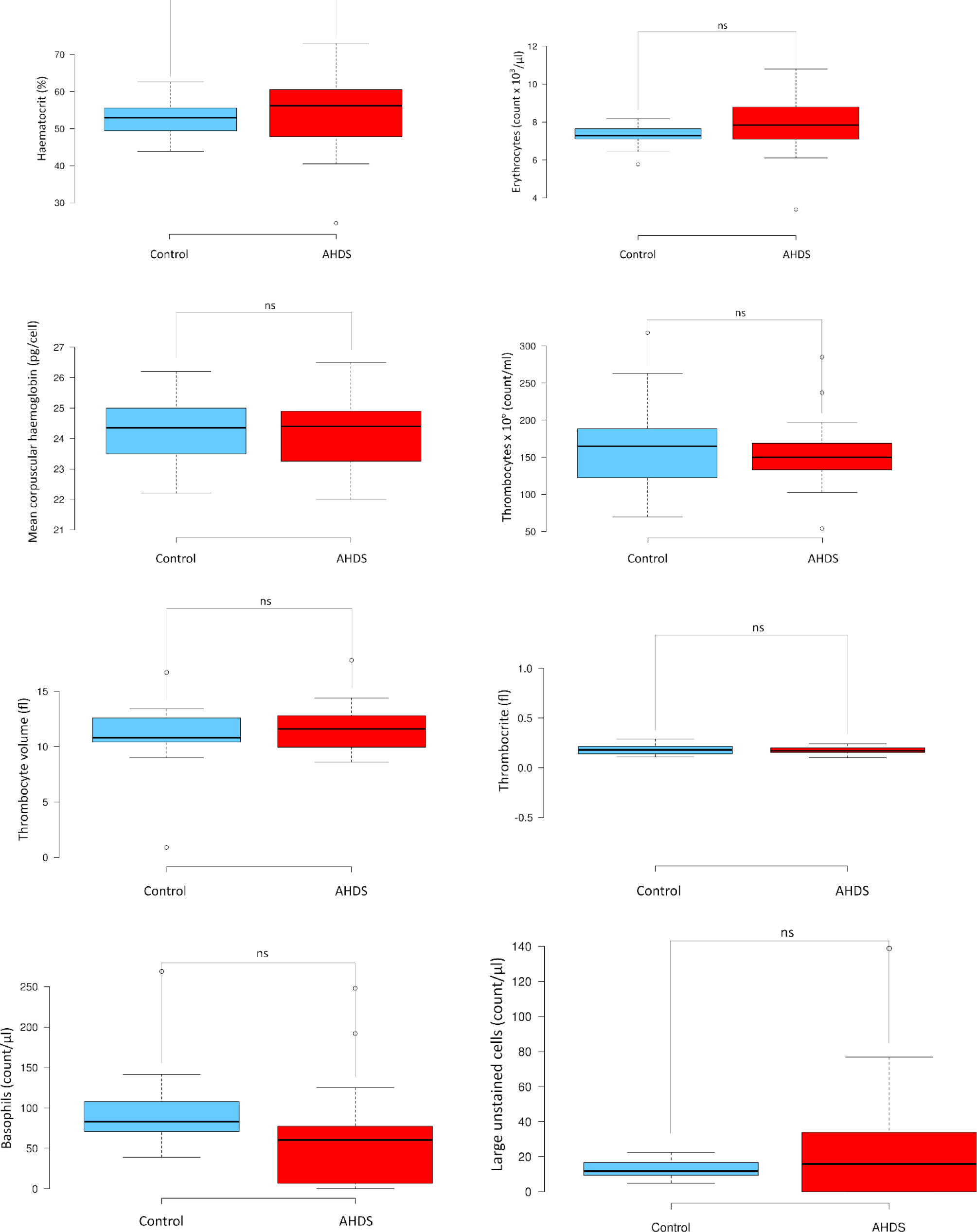
The figure displays a boxplot analysis of non-significant blood cell markers in both the AHDS and control groups. The boxplots illustrate notable variations in blood cell markers between these two groups. Statistical distinctions were assessed using the student’s t-test for samples with equal variance or the Welch test for samples with differing variances. Statistical significance was attributed to values with *p* < 0.05.

**Figure S2.**
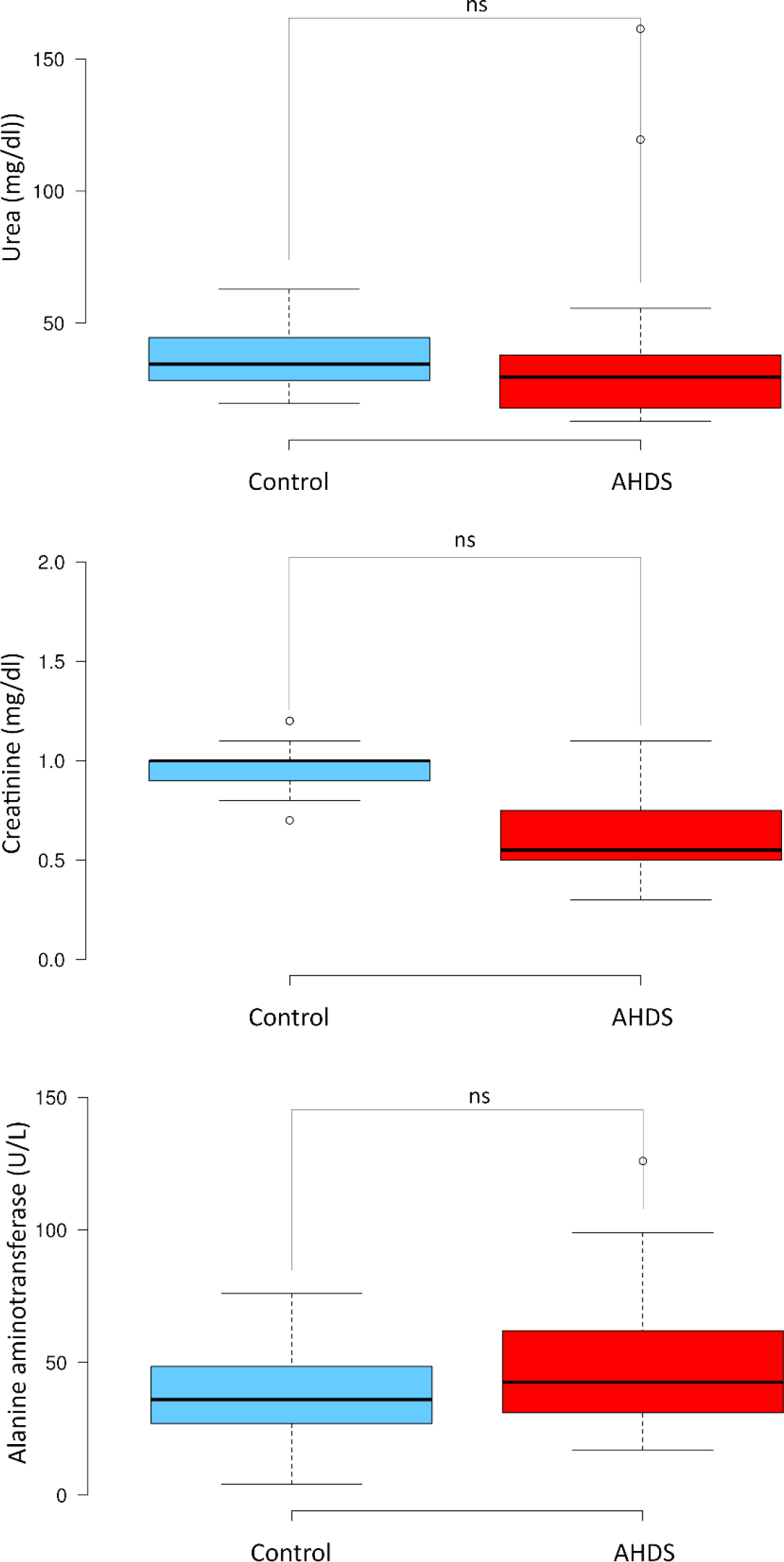
The figure illustrates a boxplot analysis of non-significant biochemistry parameters in both the AHDS and control groups. The boxplots highlight noteworthy variations in specific plasma biochemistry markers between the AHDS and control groups. Statistical differences were evaluated using either the student’s t-test for samples with equal variance or the Welch test for samples exhibiting varying variances. Statistical significance was attributed to values with *p* < 0.05.

